# Exploring non-invasive sexing of early chick embryos in intact eggs using Laser Speckle Contrast Imaging (LSCI) and Deep Neural Network (DNN)

**DOI:** 10.1101/2025.04.17.649355

**Authors:** Simon Mahler, Anika Arora, Carol Readhead, Siyuan Yin, Surya Narayanan Hari, Ellie Wang, Cecilia I. Moxley, Abdullahi A. Adeboye, Zhenyu Dong, Haowen Zhou, Xi Chen, Marianne Bronner, Changhuei Yang

**Affiliations:** Department of Electrical Engineering, California Institute of Technology, Pasadena, California 91125, USA; California Institute of Technology, Pasadena, California 91125, USA; Department of Biology and Biological Engineering, California Institute of Technology, Pasadena, California 91125, USA; Department of Medical Engineering, California Institute of Technology, Pasadena, California 91125, USA

**Author notes:** These authors contributed equally (Simon Mahler, Anika Arora, Carol Readhead, Siyuan Yin).

## Abstract

The ability to image blood flow in early-stage avian embryos has significant applications in developmental biology, drug and vaccine testing, as well as determining sex differentiation. In this project, we used our recently developed laser speckle contrast imaging (LSCI) system to non-invasively image extraembryonic blood vessels and used these images to attempt early sex identification of chick embryos. Specifically, we captured images of blood vessels from 1,251 living chicken embryos between day three and day four of incubation. We then applied deep neural network (DNN) models to evaluate whether it is possible to differentiate sex based on vascular patterns. Using ResNetBiT and YOLOv5 models, our results indicate that sex differentiation from extraembryonic blood vessel images was not achievable with sufficiently high accuracy or statistical significance for practical use. Specifically, ResNetBiT had a five-fold cross-validated average accuracy of 59%±5% (fold-wise p-value, p ≤ 0.3) at day 3 and 61%±3% (fold-wise, p ≤ 0.04) at day 4. YOLOv5 had a five-fold cross-validated average accuracy of 55%±3% (fold-wise, p ≤ 0.3) at day 3 and 53%±3% (fold-wise, p ≤ 0.5) at day 4. Our findings suggest that using vascular pattern imaging alone is inconclusive for reliable early sex identification in chicken embryos.

## Introduction

Incubating fertile chicken eggs is a crucial process in the poultry industry for producing male (cockerels) and female (hens) chicks. Hens are used for meat production and egg-laying. Cockerels can only be used for meat production, but they are generally considered economically unviable for such purpose as they grow more slowly and do not reach the same size as hens [1]. Consequently, cockerels are routinely killed immediately after sex determination (male chick culling) on the same day they hatch [2–6].

One prevalent method of culling is maceration, which involves grinding the chicks alive [4]. Other methods include gassing and cervical dislocation [5,6]. All those methods are cruel and ethically questionable due to the distress and suffering they inflict on the chicks. In addition, the newly hatched chicks have to be manually sexed which is laborious and expensive. More than seven billion male chicks are culled annually around the world [1,7]. This grim reality underscores the urgent need for reevaluation and innovation within the poultry industry to address both economic and ethical concerns associated with the treatment of male chicks.

A straightforward solution to prevent mass male chick culling is to selectively incubate only female chicken eggs. However, no effective methods currently exist to accurately identify the sex of a chicken egg before incubation [8]. An alternative approach is to incubate the eggs and apply a non-invasive sexing technology within the first few days of development. At this stage, the embryo has not yet developed a functional sensory nervous system [4,9], making it an ethical window for intervention. In addition, male chick embryos could be redirected for alternative uses such as animal research or breeding. There exist methods for early-stage sexing of avian eggs, however, they require breaking the eggshell [8,10–12] and negatively impacting the natural development of the eggs. A robust and commercially viable sexing technology should be non-invasive and should not alter egg development.

Recently, several interesting DNN-based studies have reported positive sex identification results by applying DNN methods to egg candling images of early chick embryos [8,13–15]. Egg candling is a well-established conventional method in which an egg is held up to a strong light source to reveal its blood vessel network. These studies suggest that there are subtle sex-based differences in the blood vessel network that the DNN systems can pick up.

We recently developed an improved egg imaging system using laser speckle contrast imaging (LSCI) [16,17]. Compared to traditional egg candling, this approach offers superior detection of active blood flow in blood vessels and is robust against eggshell coloration variations and stains [16]. We anticipate that blood vessel images acquired with our LSCI system would serve as better input for DNN-based image analysis. In this project, we aim to objectively evaluate whether DNN can be used conclusively to determine sex for early-stage chick embryos in intact eggs.

In this paper, we applied our non-invasive LSCI system to capture images of the active extraembryonic blood vessels of chicken embryos between day 3 and day 4 of incubation. Blood vessel images were recorded from 1,251 chicken eggs. The eggs were then re-incubated until day 11, at which point the sex was confirmed by breaking the eggs to directly examine the gonads and confirm the sex genetically by PCR. Using the collected dataset, we trained two deep neural network (DNN) models (ResNetBiT and YOLOv5) for the task of identifying the sex of the chick eggs based on blood vessel images. Our results, from a five-fold cross-validation using ResNetBiT or YOLOv5 DNNs, suggest that the model may have some capabilities to differentiate between male and female LSCI images but not at a high enough level for practical application. ResNetBiT had an average accuracy of 59% (fold-wise p-value, p ≤ 0.3) at day 3 and 61% (fold-wise, p ≤ 0.04) at day 4. YOLOv5, had an average accuracy of 55% (fold-wise, p ≤ 0.3) at day 3 and 53% (fold-wise, p ≤ 0.5) at day 4. These results are in disagreement with those obtained using a similar protocol in Ref. [13], where an accuracy of around 85% was achieved with egg candling images of comparable dataset size, and similar algorithm (YOLOv5). We conclude by pointing out further avenues of study.

## 1. Materials and Methods

### 1.1 Laser Speckle Contrast Imaging (LSCI) of chick embryo blood vessels

Figure 1 presents a typical image acquired with our laser speckle contrast imaging (LSCI) system [16–18], which operates based on laser speckles generated by the scattering of coherent laser light within the sample [18–24]. A laser speckle pattern is characterized by a random granular distribution of bright and dark spots. In our case, the laser speckles result from the interference of randomly scattered laser light by the eggshell and various internal components of the egg, including red blood cells flowing through the blood vessels, the yolk, the embryo, and the albumen [16,18,25– 27]. See Refs. [16,17] for more details on LSCI applied to image the chick embryo blood vessels.

**Figure 1:**
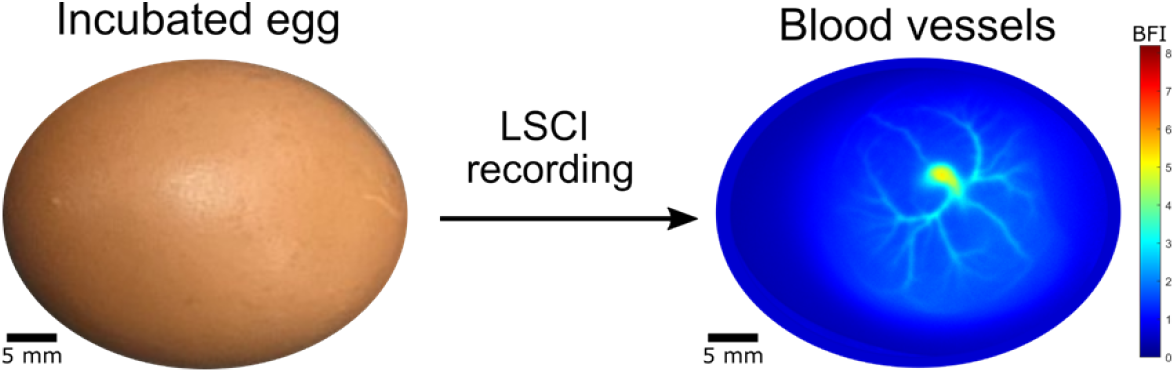
Representative LSCI image of flowing blood vessels in a chick embryo, obtained non-invasively through the eggshell. See online version for best visualization of the blood vessel image.

As components such as blood move within the egg, the speckle field undergoes temporal changes. Since blood flows at a higher speed compared to other components of the egg, such as the yolk or albumen, the temporal speckle variations are primarily influenced by blood movement. In our case, LSCI is accomplished by illuminating the egg with coherent laser light from the side of the egg and collecting the scattered light from the top of the egg with a high-resolution camera [16,17].

Speckle contrast, which typically ranges from 0 to 1, quantifies intensity variations within a speckle pattern. As blood cells move within blood vessels, their motion generates multiple speckle realizations during the camera’s exposure time. When these fluctuations occur faster than the integration time of a single image frame, they are averaged out, leading to a smeared and washed-out speckle pattern recorded by the camera. In contrast, when fluctuations are much slower than the integration time, the speckle pattern remains more defined, and pixel intensity readings reflect greater variation. Therefore, lower speckle contrast corresponds to regions with faster motion, such as flowing blood, whereas higher speckle contrast indicates static or slow-moving regions. Speckle contrast is mathematically defined as the ratio of the standard deviation to the mean value of pixel intensities, Eq. (1).

In our LSCI system (Appendix Fig. A1), the laser light source was a single-frequency continuous-wave 852 nm laser [Spectra-Physics DL852-300-SO] with an output power of 230 mW. The imaging setup included a 12.3-megapixel CMOS camera [Thorlabs CS126CU] with a pixel size of 3.45 × 3.45 µm^2^, operating at a frame rate of 21 frames per second and a pixel bit-depth of 12 bits. The camera exposure time was set to 10 ms. A 50 mm focal length lens [Edmund Optics #86-574] together with the camera formed an imaging system, achieving a magnification ratio of 0.2 and a numerical aperture of 0.03. This configuration resulted in a speckle field at the camera where, on average, a speckle would span 2 pixels laterally. The camera’s quantum efficiency at the laser wavelength of 852 nm was approximately 20%. In future studies, we plan to utilize a camera with higher quantum efficiency and a faster frame rate. See Refs. [16,17] for more details on the experimental arrangement. Note that our LSCI system can also monitor blood flow time trace, with similar temporal resolution and sensitivity as speckle contrast optical spectroscopy (SCOS) [16,28,29].

Since the movement of blood flow within the vessels is significantly greater than that of other egg components, a distinct contrast difference emerges between the blood vessels and surrounding structures. Specifically, the higher motion of blood cells leads to lower speckle contrast, whereas areas with less motion exhibit higher speckle contrast. This presents a key distinction from the traditional egg candling method, which relies on brightfield imaging to measure differences in light absorption between blood vessels and other components [8,13,16]. Unlike brightfield imaging, which detects absorption contrasts, LSCI selectively differentiates dynamic components from static ones [16]. Consequently, if a blood vessel is inactive (i.e., without flowing blood), it would not appear in LSCI imaging but would still be visible in brightfield imaging. This fundamental difference underscores the unique capability of LSCI in assessing blood flow dynamics. A detailed comparison study between the two methods can be found in [16].

For this study, we used temporal LSCI [30,31], where the camera recorded a sequence of N = 100 speckle frames images when the laser was on and another 100 noise frames *Ĩ*(*t*) when the laser was blocked. Both sets had the same exposure time T. The noise frames were averaged into a single frame and subtracted from the speckle frames [16]. The temporal speckle contrast *K*_*t*_ was calculated for each camera pixel *Ĩ*(*i,j;t*) in the temporal domain over the N noise-subtracted speckle frames as [16,17]:

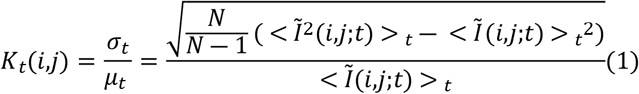

where *Ĩ*(*i,j*) is the intensity at the pixel row *i* and column *j, t* is the time at which the speckle pattern was recorded by the camera, and <> _*t*_ indicates temporal averaging over time occurring at pixel *i,j*. The speckle contrast corresponds to the ratio between the standard deviation *σ*_*t*_ and the temporal mean *μ*_*t*_ of the N recorded speckle pattern images. The relative blood flow index (BFI) was estimated from the speckle contrast as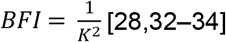. Note that this estimation is valid when the exposure time of the camera is much longer than the speckle field decorrelation time and rely on the assumption that the motion of multiple scattering blood cells is unordered [33]. It also assumes that the scattered light is an ergodic field with the absence of static scattering. These considerations are extensively studied in Ref. [35], as the different scattering regimes and particle motion types can change the form of the field correlation function *g*_1_(*τ*) [33,35,36]. See Appendix A of Ref. [16] and Ref. [33] for more analysis on the blood flow index calculation from speckle contrast images in our LSCI system. The blood flow index provides relative information on blood flow, accounting for the total volume of blood moved within a given time period, differing from blood volume [37,38]. Figure 1 shows a typical example of an egg’s blood vessel image with a clear visualization of the blood vessel network. Our method can image blood vessels as small as 100 µm, see Ref. [16].

### 1.2 Egg incubation

For this study, brown fertile eggs were obtained from Sun Valley Farms and Petaluma Poultry in California. The eggs were incubated using Hethya, HHD, and Kebonnixs commercial incubators, which automatically maintained the temperature inside the incubator between 37–38.5°C (99–101°F) and gently rotated the eggs every 60 to 90 minutes. Humidity levels were kept within a 50–75% range. After three days of incubation, embryos that had developed beyond stage HH17 were selected for imaging. All animal research procedures were approved by the Caltech Office of Laboratory Animal Resources. No eggs were incubated beyond day 12, and embryo disposal was conducted in compliance with the Caltech Institutional Animal Care and Use Committee (IACUC) policies.

### 1.3 Speckle imaging data collection protocol and ground-truth sex

Our data collection protocol was as follows: first, fertile eggs were incubated in a large incubator capable of holding up to 60 chicken eggs. Each egg was assigned a unique number between 1 and 1,500 in chronological order (i.e., the first egg was labeled 1, the second 2, and so on). The label was written with a black permanent marker on the rounded bottom part of the egg. The incubator automatically maintained the internal temperature and gently rotated the eggs. Distilled water was added whenever the humidity level dropped below 55%.

Second, after three days of incubation, we performed LSCI imaging (as in Fig. 1) at different developmental Hamburger and Hamilton (HH) [40] stages of HH18 (68 hours of incubation), HH19 (72 hours), and HH20 (75 hours) (Fig. 2). After each imaging session, the egg was returned to the incubator. To prevent the same egg from being used in both training and validation within a single DNN experiment, LSCI images from the same egg were used exclusively for either training or validation.

**Figure 2:**
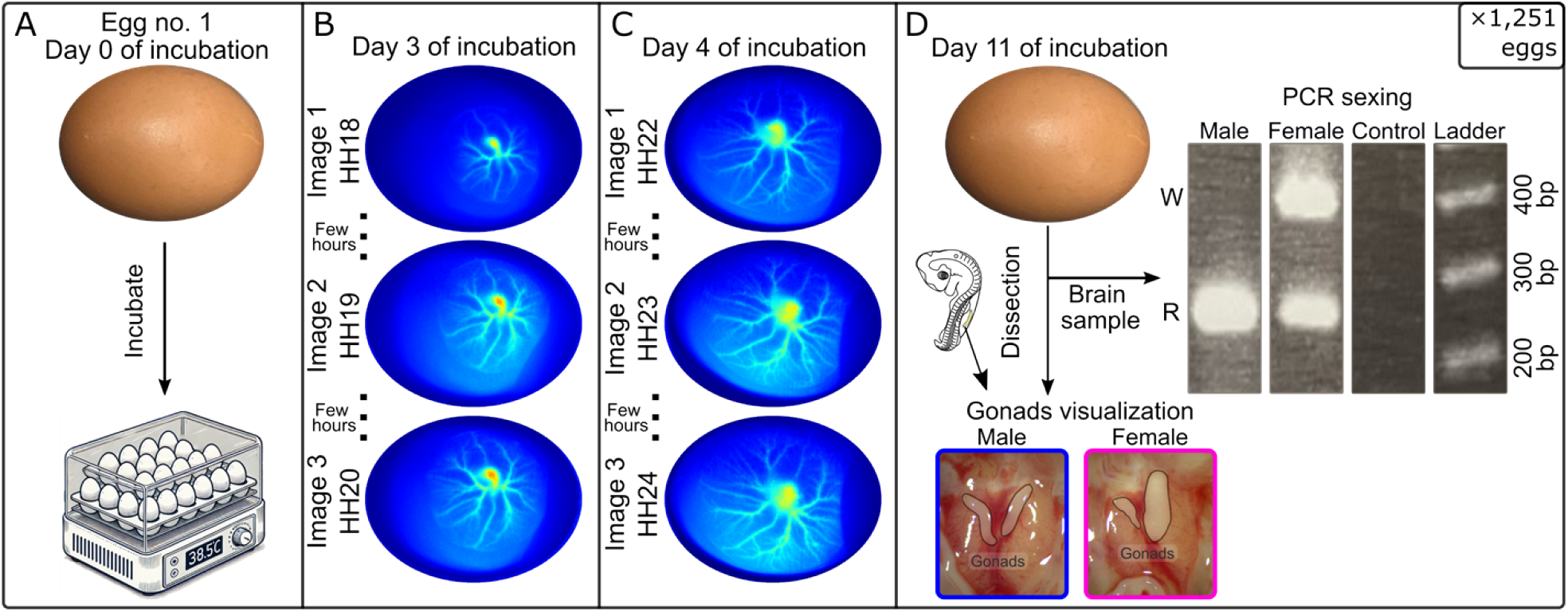
Speckle imaging data collection protocol for a single egg. (A) A fertile egg (e.g., Egg 1) was incubated in a large incubator. (B) After three days of incubation, the egg was imaged using our LSCI system. (C) After four days of incubation, the egg was imaged again using our LSCI system. (D) After eleven days of incubation, the eggshell was broken, and the developing chick embryo was dissected to visualize the gonads for sex identification. At the same time, a brain sample was collected and frozen at - 20^0^C for PCR genetic testing to confirm the visual sex identification. Male and female examples are shown for comparison. See online version for best visualization of the blood vessel images.

Third, after four days of incubation, each egg was LSCI imaged at different developmental stages of HH22 (92 hours), HH23 (96 hours), and HH24 (100 hours) (Fig. 2). If the imaging quality was insufficient to distinguish blood vessels from the background, the LSCI images were discarded. Due to time constraints and imaging quality selection, most eggs were only imaged at one or two developmental stages. After imaging, the eggs were returned to the incubator until day 11.

Lastly, after eleven days of incubation, the eggshell was broken, and the chick embryo was carefully dissected so that the gonads could be visualized for sex identification [39,40]. An incision was made on the ventral surface of the embryo, and the gonads were exposed (see Fig. 2D insets). In males, the left and right gonads were typically similar in size and were thin, elongated structures [39,40]. In females, the left gonad appeared larger and more elongated than the right gonad. The images were recorded using a 0.7X-5X magnification ratio microscope (AmScope H800 Series). The sex of the embryo was further validated by using PCR [41]. For the PCR analysis, approximately 50 mg of soft brain tissue was collected at the time of dissection and stored at -20°C. (Fig. 2D insets). The tissue was extracted using Thermo Scientific™ Animal Tissue Direct PCRKit according to the manufacturer’s instructions. The primers for the PCR reaction were the W-repeat (W), which is specific to female DNA and R the 18S ribosomal gene which is present in both sexes Ref. [41]. The primers used were W from the female specific W chromosome which was designed to amplify a 415 Bp product, and R from the 18S ribosomal gene, which were designed to amplify a 256 Bp product Ref. [41].

The primers were:

W 5’ s primer: 5′ CCCAAATATAACACGCTTCACT 3′,

W 3’ s primer: 5′ GAAATGAATTATTTTCTGGCGAC 3′,

R 5’ s primer: 5′ AGCTCTTTCTCGATTCCGTG 3′,

R 3’ s primer: 3′ GGGTAGACACAAGCTGAGCC 3′.

Figure 2D presents a typical visualization of gonads alongside PCR results for both sexes. The PCR products were run on a 1.5% agarose gel. In females, both the W and R products are detected at 415 and 256 bp, respectively, whereas in males, only the R sequence is present. On average, there was a 2% discrepancy between the sex identification via PCR and gonadal visualization. LSCI images from these eggs were discarded to maintain consistency within the dataset.

### 1.4 Dataset Overview: LSCI Images and Egg Count

Following the LSCI imaging protocol (Fig. 2), we recorded a total of 2,773 LSCI images collected from 1,251 eggs. Table 1 summarizes the distribution of images and eggs across incubation days and sex categories. On day 3, covering developmental Hamburger and Hamilton (HH) [42] stages HH18 to HH20, a total of 1,552 images were collected, with 52% from male and 48% from female embryos. On day 4, spanning HH22 to HH24, 1,220 images were collected, with a similar distribution between males (52%) and females (48%). Both days combined, 2,772 images were captured, with 51% from male embryos and 49% from female embryos. Readers interested in viewing the LSCI images are invited to see Appendix figures B1 and B2, which display 16 LSCI images randomly selected from day 3 and day 4 datasets.

**Table 1:** Statistical summary of the LSCI images dataset with female and male ratios. A total of 2,773 images were collected from 1,251 eggs.

| Incubation Day | Sex | No. of Eggs | No. of Images | Total No. of Images |
| --- | --- | --- | --- | --- |
| Day 3<br>(HH17-HH20) | Male | 598 (52%) | 794 (51%) | 1,553 |
|  | Female | 543 (48%) | 759 (49%) |  |
| Day 4<br>(HH21-HH25) | Male | 479 (52%) | 631 (52%) | 1,220 |
|  | Female | 440 (48%) | 589 (48%) |  |
| Days 3 & 4 | Male | 652 (48%) | 1,425 (51%) | 2,773 |
|  | Female | 599 (52%) | 1,348 (49%) |  |

The number of images collected per egg was not uniform due to time constraints, as we can only image the eggs during relatively narrow time windows. In addition, some images had to be rejected due to poor imaging conditions, such as bad embryo positioning, embryonic body movements, or artifact noise. However, these experimental variations have little impact on model performance, given that we utilized transfer learning with models pretrained on ImageNet, a large-scale dataset containing over 14 million labeled images across thousands of diverse categories [43]. Transfer learning from such large-scale datasets typically provides models with robustness against minor class imbalances in downstream tasks [44,45]. For validation, we selected one image per egg to ensure balanced representation across samples and to prevent any bias that could arise from over-representing individual embryos. This strategy prevents potential bias from over-representation of individual embryos, ensures statistical robustness, and allows for a fair evaluation of the model’s generalizability.

### 1.5 Data Preparation and Preprocessing

We followed a standardized approach for preprocessing the LSCI images, shown in Figure 3. We used monochrome grayscale images as input for the DNNs, on the assumption the vascular structure could serve as the primary indicator of sex. First, we applied a center crop to the original image, reducing its dimensions from 4,096 × 3,000 pixels to 3000 × 3000 pixels. To enhance vascular structure, we implemented a sliding window technique to detect the brightest 30 × 30 pixels region, identifying it as the embryonic heart. We then zoomed into this region by cropping a 2,300 x 2,300 pixel square centered around the brightest area. To ensure compatibility with pretrained DNNs, we resized all images to 256 × 256 pixels. Additionally, we applied various image enhancement techniques to optimize feature extraction. Unsharp masking [46,47] was used for image sharpening to enhance image details by subtracting a blurred version of the image from the original. The Scharr operator [48] was employed for edge detection to emphasize gradient edges and features. Finally, Contrast Limited Adaptive Histogram Equalization (CLAHE) [49,50] was applied to enhance contrast while preventing over-amplification of noise.

**Figure 3:**
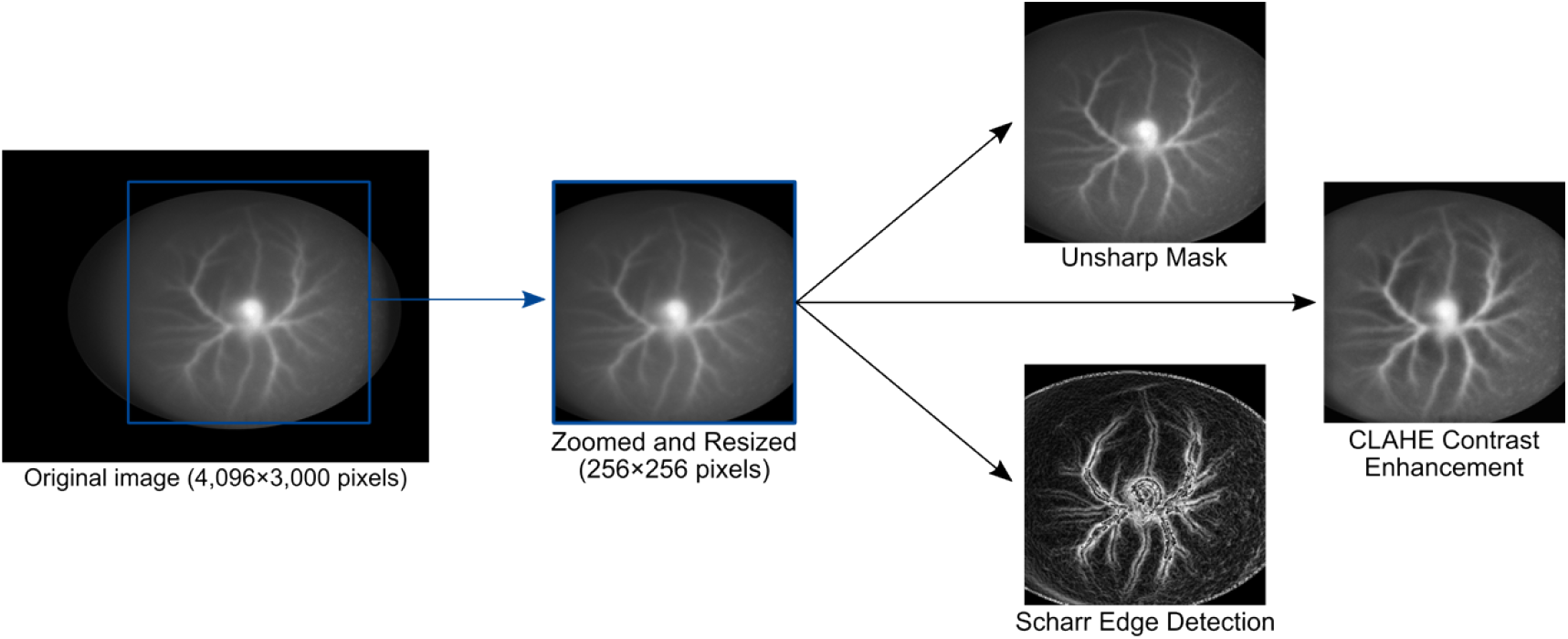
Chicken embryo dataset pre-processing pipeline. Each LSCI image undergoes the same pre-processing pipeline before being input into the algorithm.

We split the complete dataset from days 3 and 4 (Table 1) into training and validation sets with an 80/20 split ratio. For the validation set, we used one image per egg. Data augmentation was applied exclusively to the training set to improve model generalization and prevent overfitting. The data augmentations included random rotations up to 180°, translations up to 5% in any direction, image scaling between 0.85x and 1.15x, and random vertical and horizontal flips with probability 0.3. No augmentations were applied to the validation set. We implemented five-fold cross-validation to evaluate the model’s performance.

### 1.6 Choice of the Deep Neural Networks (DNNs)

We initially trained and tested four DNNs, including ResNetBiT (Big Transfer ResNet) [51], Xception41 [52], DenseNet121 [53], and InceptionV3 [54]. This preliminary assessment was done using a train/test split instead of cross-validation. Model performance was evaluated based on accuracy, area under the receiver operating characteristic curve (AUC), and loss. The AUC, a standard metric for assessing a model’s ability to distinguish between classes (here, female vs. male), ranges from 0 to 1, with higher values indicating better balance between sensitivity (true positive rates for females) and specificity (true negative rates for males). Accuracy measures the proportion of correctly classified samples, while loss quantifies the error between the predicted and actual labels, with lower values indicating better model performance.

When training accuracy and AUC are high on training data but significantly lower on validation data, and when training loss decreases while the loss stagnates or increases, it indicates overfitting [55]—where the model memorizes training data but fails to generalize to unseen data. The binary cross-entropy (BCE) loss function penalizes incorrect predictions more heavily than it rewards correct ones, making it an effective tool for identifying overfitting [56,57]. Therefore, we applied BCE loss in all the experiments presented in this paper. The BCE loss *L* for a set of *n* predictions (*ŷ*_1_, *ŷ*_*2*_, …, *ŷ*_*n*_), given true labels (*y*_1_, *y*_*2*_, …, *y*_*n*_) is as follows:

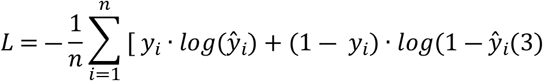

After training and validating the four DNNs for 25 epochs, the accuracy, AUC, and loss for each model were as follows: ResNetBiT (accuracy: 51%, AUC: 0.51, loss: 0.72), Xception41 (accuracy: 54%, AUC: 0.54, loss: 0.80), DenseNet121 (accuracy: 53%, AUC: 0.54, loss: 0.85), and InceptionV3 (accuracy: 55%, AUC: 0.57, loss: 0.76). Although ResNetBiT performed slightly worse overall, it exhibited the least overfitting with lowest loss and with training and test curves showing similar trends (not shown). This suggests that ResNetBiT may have greater learning potential if trained for additional epochs. Therefore, we shifted our focus to optimizing ResNetBiT performance.

In addition to ResNetBiT, we also tested a YOLO-based model, as Ref. [13] reported its performance in sex identification of chicken embryos using blood vessel images obtained from egg candling. In that study, a slightly modified YOLOv7 model incorporating a convolutional block attention module (CBAM) achieved an accuracy of 85% or higher on incubation days 3 and 4, with at least 87% precision, 82% recall, and 85% average precision. Reference [13] also compared four models—Faster R-CNN, SSD, YOLOv5, and their modified YOLOv7—for sex identification from blood vessel images on day 4 of incubation. Reference [13] used a dataset comprised of 5,940 images from 2,844 eggs, with an approximate 70/30 training-to-testing split. The results indicated that YOLO models were more effective at identifying blood vessel regions and differentiating sex, with YOLOv5 and modified YOLOv7 exhibiting less than 3% deviation in precision and recall. Consequently, both models are expected to achieve an accuracy of at least 85%. Since we did not have access to the modified YOLOv7 model from Ref. [13], we instead used the standard YOLOv5 model as a comparative baseline for evaluating our results against those reported in the study.

### 1.7 Approaches: ResNetBiT and YOLOv5

To classify the sex of chicken embryos, we used two DNN architectures, ResNetBiT [51] and YOLOv5 [58], both adapted for binary classification. ResNetBiT is a high-capacity convolutional neural network designed for robust feature extraction and transfer learning. Pretrained on the ImageNet-21k dataset, it is well-suited for transfer learning on small datasets. We initialized ResNetBiT with pretrained weights and fine-tuned it on our dataset. Similarly, we adapted YOLOv5, a model typically used for object detection, for classification. This approach allowed us to explore whether a detection-based model, originally designed for localization tasks, could better suit this problem. For the YOLOv5 model, we trained on 640 x 640 pixel images to match the default image size for the model.

Both models were trained using the Adam optimizer, BCE loss function, and with the Cosine Annealing learning rate scheduler to ensure stable convergence. To prevent overfitting, early stopping was applied if the accuracy did not increase for ten consecutive epochs, following a minimum number of training epochs (40 epochs for ResNetBiT, and 15 epochs for YOLOv5). The ResNetBiT model was run for a maximum of 100 epochs, while YOLOv5 was trained for up to 300 epochs. We trained on one-channel grayscale images using each of the preprocessing techniques of Fig. 3— normal images, unsharp masking, edge detection, and contrast enhancement.

## 2. Results

From the days 3 and 4 dataset, we conducted a final five-fold cross-validation run with trained ResNetBiT and YOLOv5 models. For each fold, the training set contained approximately 2,200 images, while the validation set contained around 230 images for day 3 dataset, and 185 images for day 4 dataset. The exact sizes varied slightly between folds because the splits were designed to ensure that images from the same egg were included exclusively in either the training or validation set, and that only one image per egg was included in the validation set.

### 2.1 ResNetBiT

The five-fold cross-validation results for the ResNetBiT model are presented in Fig. 4 and Table 2. Figure 4A depicts the training and validation accuracy curves over the course of epoch. The validation curves are averaged over the five folds. As shown, the training accuracy increases significantly with the number of epochs, surpassing 75% after 50 epochs. However, the validation accuracy curves for day 3 and day 4 datasets fluctuates between 50% and 63%, with average accuracies across five-fold runs of 59% for the day 3 dataset and 61% for the day 4 dataset after 50 epochs. The p-value for each fold was calculated using a binomial calculator assuming a random guess [59]. We establish the following significance levels α: p > 0.05 (ns: not significant), p < 0.05 (*), p < 0.01 (**), p < 0.001 (***), and p < 0.0001 (****).

**Figure 4:**
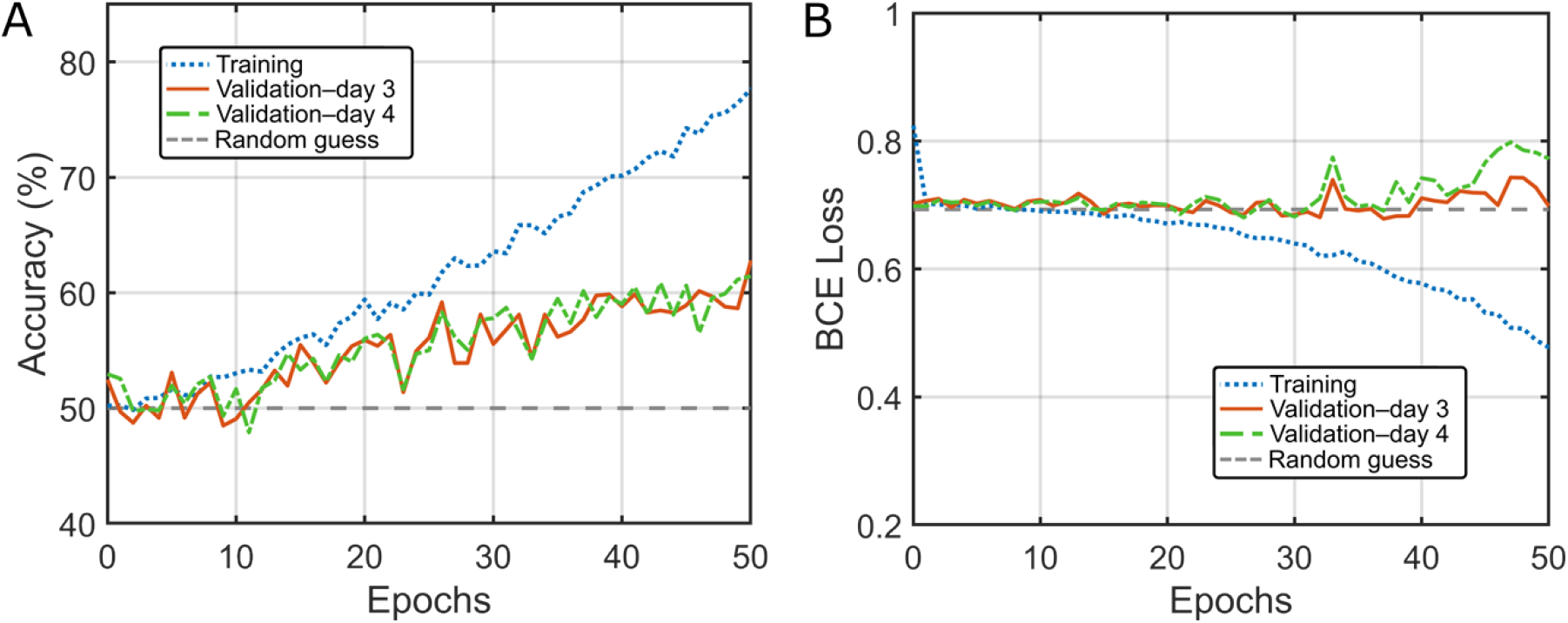
Sex identification results on LSCI images using the ResNetBiT model. (B) Average accuracy and (C) BCE loss for the training (blue dotted curve) and validation sets on day 3 (line orange curve) and day 4 (dashed line green curve) datasets as functions of the number of epochs.

**Table 2:** Five-fold cross-validated accuracies and p-value results for the ResNetBiT model at day 3 and day 4 of incubation. The significance levels are: p > 0.05 (ns: not significant), p < 0.05 (*), p < 0.01 (**), p < 0.001 (***), and p < 0.0001 (****).

| ResNetBiT | Day 3 (HH17-HH20) |  |  | Day 4 (HH21-HH25) |  |  |
| --- | --- | --- | --- | --- | --- | --- |
| | Accuracy | p-value | $\alpha$ | Accuracy | p-value | $\alpha$ |
| Fold 1 | 62% | 0.0006 | *** | 61% | 0.002 | ** |
| Fold 2 | 58% | 0.014 | * | 62% | 0.001 | ** |
| Fold 3 | 61% | 0.0004 | ** | 57% | 0.04 | * |
| Fold 4 | 61% | 0.0007 | ** | 64% | 0.0001 | ** |
| Fold 5 | 52% | 0.3 | ns | 61% | 0.001 | ** |

Table 2 shows the accuracy, p-value, and significance level α for each fold of the ResNetBiT model tested on datasets from day 3 and day 4 of incubation. As the five-fold experiments are performed with the same dataset, their prediction accuracy and associated p-value are not independent of each other. While the accuracy averaged across the five-fold provide some measure of the overall accuracy, it would be meaningless to attempt averaging the p-value. A more meaningful approach would be to look at the worst performing p-value across the folds. Doing so, we can see, from Table 2, that the worst performing p for ResNetBiT for the day 3 experiment was 0.3. For day 4 experiment, the worst performing p was 0.04. Using our established significance level, only ResNetBiT day 4 model appeared to barely pass the threshold for *. The low averaged accuracy of 60% is also noteworthy in that it is not a high enough accuracy for practical applications. These findings are further supported by the training and validation BCE loss curves in Fig. 4B. The training loss decreases from 0.7 to around 0.45 over 50 epochs, while the validation loss curves slightly increase from 0.70 to 0.80 over 50 epochs. This suggests overfitting of the model, confirming that it is not suitable for practical application. For a random binary guess, the expected BCE loss is *log*(*2*) ≈ 0.693.

### 2.2 YOLOv5

The five-fold cross-validated results for the YOLOv5 model are presented in Fig. 5 and Table 3. Figure 4A depicts the training and validation accuracy curves over the course of epochs. The validation curves are averaged over the five folds. As shown, the training accuracy increases significantly with the number of epochs, approaching 100% after 30 epochs. However, the validation accuracy curves for day 3 and day 4 datasets fluctuate between 50% and 58%, with average accuracies across five-fold runs of 55% for the day 3 dataset and 53% for the day 4 dataset. The worst performing p for YOLOv5 for the day 3 experiment was 0.3. For day 4 experiment, the worst performing p was 0.5. Using our established significance level, neither passed the significance threshold. These findings are further supported by the training and validation BCE loss curves in Fig. 4B. The training loss decreases from 0.7 to around 0.2 over 30 epochs, while the validation loss curves slightly increase from 0.70 to 0.90 over 30 epochs, suggesting overfitting. For a random binary guess, the expected BCE loss is *log*(*2*) ≈ 0.693. Table 3 shows the accuracy, p-value, and significance level α for each fold of the ResNetBiT model tested on datasets from day 3 and day 4 of incubation.

**Figure 5:**
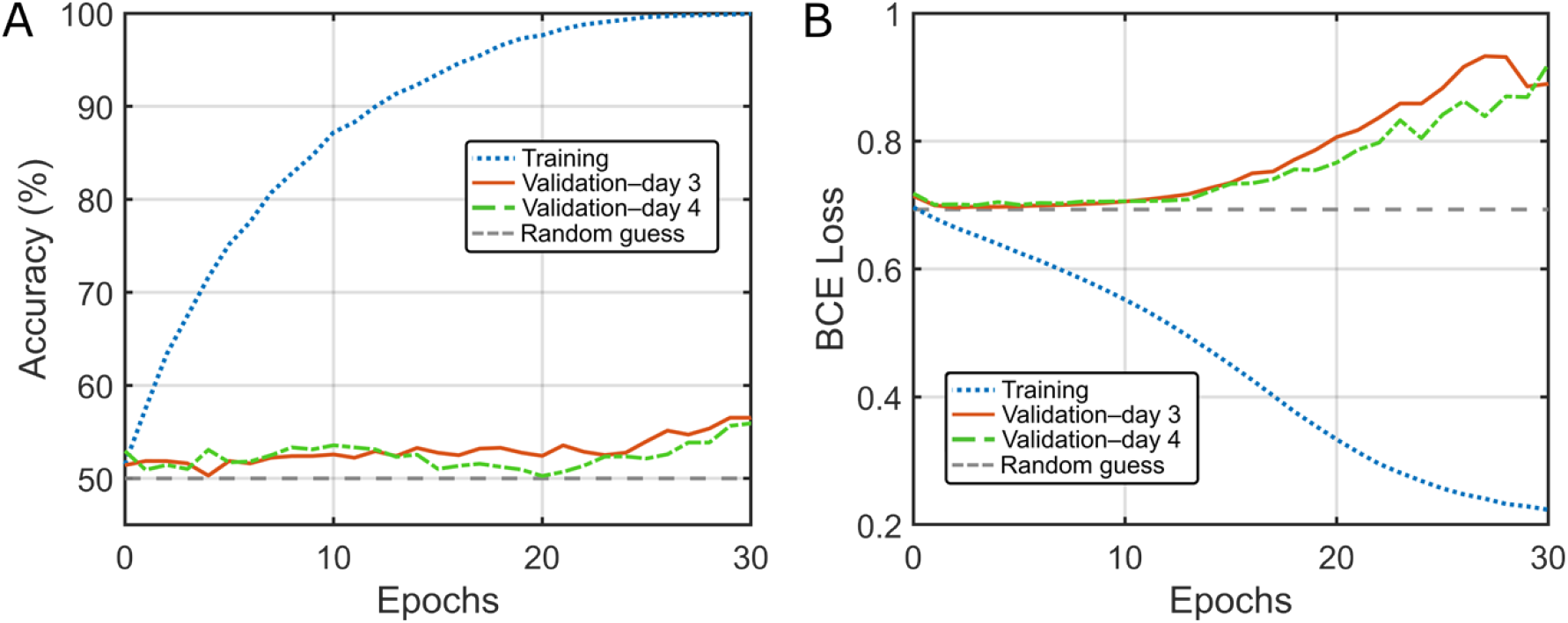
Sex identification results on LSCI images using the YOLOv5 model. (B) Average accuracy and (C) BCE loss for the training (blue dotted curve) and validation sets on day 3 (line orange curve) and day 4 (dashed line green curve) datasets as functions of the number of epochs.

**Table 3:** Five-fold cross-validated accuracies and p-value results for the YOLOv5 model at day 3 and day 4 of incubation. The significance levels are: p > 0.05 (ns: not significant), p < 0.05 (*), p < 0.01 (**), p < 0.001 (***), and p < 0.0001 (****).

| YOLOv5 | Day 3 (HH17-HH20) |  |  | Day 4 (HH21-HH25) |  |  |
| --- | --- | --- | --- | --- | --- | --- |
| | Accuracy | p-value | $\alpha$ | Accuracy | p-value | $\alpha$ |
| Fold 1 | 58% | 0.014 | * | 53% | 0.2 | ns |
| Fold 2 | 52% | 0.3 | ns | 50% | 0.5 | ns |
| Fold 3 | 54% | 0.1 | ns | 56% | 0.07 | ns |
| Fold 4 | 57% | 0.02 | * | 57% | 0.04 | * |
| Fold 5 | 53% | 0.2 | ns | 52% | 0.3 | ns |

Additionally, the classification performance was independently validated using ResNet50 and YOLOv8. The system processes egg images from both day 3 (HH19) and day 4 (HH25) developmental stages, with capabilities to handle either individual stage or a mixed dataset approach. Two complementary neural network architectures were implemented: a modified ResNet50 model adapted for grayscale image processing with an option for transfer learning from ImageNet weights, and a YOLOv8 classification model leveraging recent advancements in object detection architectures. Both models were trained with optimization for binary sex classification, incorporating data augmentation, normalization techniques, and performance monitoring through Weights & Biases integration. Experimental results demonstrated similar results around random performance in sex identification, with model performance metrics tracked across multiple training configurations.

### 2.3 Comparison with Previous Studies

Our study was unable to accurately distinguish the sex of chick embryos from LSCI images at days 3 and 4. We trained two DNNs, ResNetBiT and YOLOv5, neither achieve practically useful accuracy (80% or better). In fact, none achieved statistical significance, except ResNetBiT for day 4 images which barely pass the significance level of *.

These results are puzzling given that Ref. [13] reported over 85% accuracy with a low loss using YOLO models on chick embryo blood vessel images from day 3 or 4 of incubation. Additionally, Ref. [13] compared YOLOv5 and their modified YOLOv7 for sex identification from blood vessel images on day 4 of incubation, using a dataset of 5,940 images from 2,844 eggs with a 70/30 train-test split. Their results demonstrated that YOLO models effectively identified blood vessel regions and differentiated sex, with YOLOv5 and modified YOLOv7 showing less than a 3% deviation in precision and recall. Based on these findings, both models were expected to achieve at least 85% accuracy, making our lower performance particularly unexpected. We could not pinpoint a major factor explaining the disparity between results. Their training set was 1.9 times as large, but we do not expect this alone would account for such a significant deviation in accuracy. Despite multiple attempts, we were unable to obtain access to the dataset and/or code used in Ref. [13].

## Conclusion

Using laser speckle contrast imaging (LSCI), we recorded images of chick embryo blood vessels at days 3 and 4 of incubation to explore the feasibility of sex differentiation using deep neural network (DNN) models. Our final dataset consisted of images from 1,251 chick embryos, providing a substantial sample size for a pilot DNN testing. We trained two DNNs, ResNetBiT and YOLOv5, both of achieved not satisfactory accuracy and relatively high BCE loss, with YOLOv5 performing worse than ResNetBiT. A per-egg evaluation approach did not provide high accuracy or compelling statistical significance, leading us to conclude that sex identification from LSCI images at this early incubation stage is likely not feasible for practical applications.

This finding aligns with previous literature on genetic expression related to sex differentiation in embryos, which indicates that most genes influencing sex determination begin to express around day 4.5 of incubation in male embryos and day 6.5 in female embryos [60]. From day 6.5 onward, gradual morphological changes in gonadal tissue become observable, marking the onset of sex organ development. Additionally, the chorioallantoic membrane (CAM)—a highly vascularized extra-embryonic membrane—plays a crucial role in embryonic development, supporting vital functions such as gas exchange, calcium absorption, and waste removal. Since the CAM begins to form around day 5 of incubation (stage HH25) and is closely linked to the embryo’s genetic identity and overall development, we hypothesize that it may also be involved in sex differentiation. Given that the CAM is rich in active blood vessels and continues developing until days 12–14, this could explain why sex identification through blood vessel imaging may become more viable at later incubation stages. While we hypothesize that sex differentiation might be detectable from day 6 onward, our LSCI imaging system is currently limited to day 5.5, as increasing embryo size and movement beyond this stage disrupts image acquisition.

Future studies could explore alternative approaches, such as multiple-instance learning to assess spatial and temporal variations in blood vessel patterns, feature extraction from LSCI images, blood flow dynamics and heart rate analysis, which would require precise temperature control during image acquisition for accurate measurements. These refinements may offer new insights into the potential for early-stage sex differentiation in chick embryos using LSCI imaging.

## Appendix

### Appendix A: Experimental arrangement of the LSCI system

**Figure A1:**
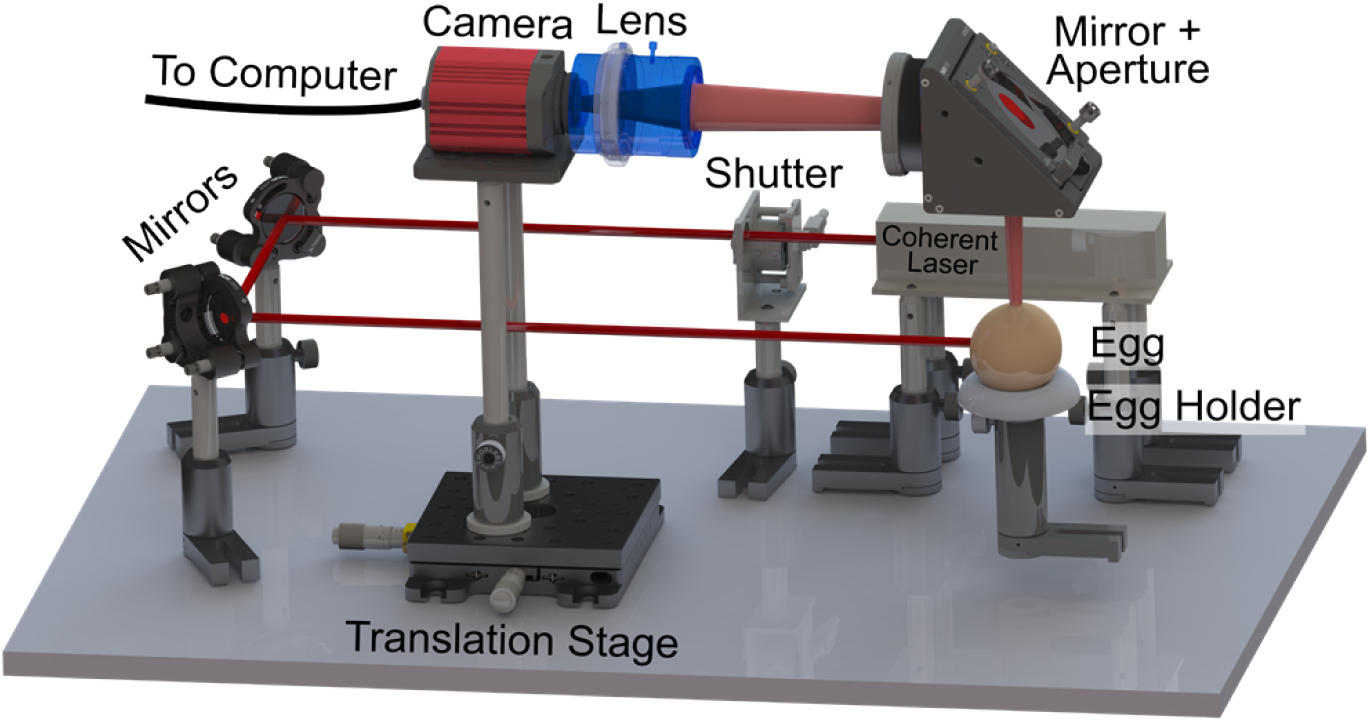
Experimental arrangement of our LSCI system for imaging the active blood vessels of a chick embryo non-invasively. The whole system was encased in a black box to avoid stray light. See Ref. [16] for a detailed experimental arrangement.

### Appendix B: Examples of LSCI images

**Figure B1:**
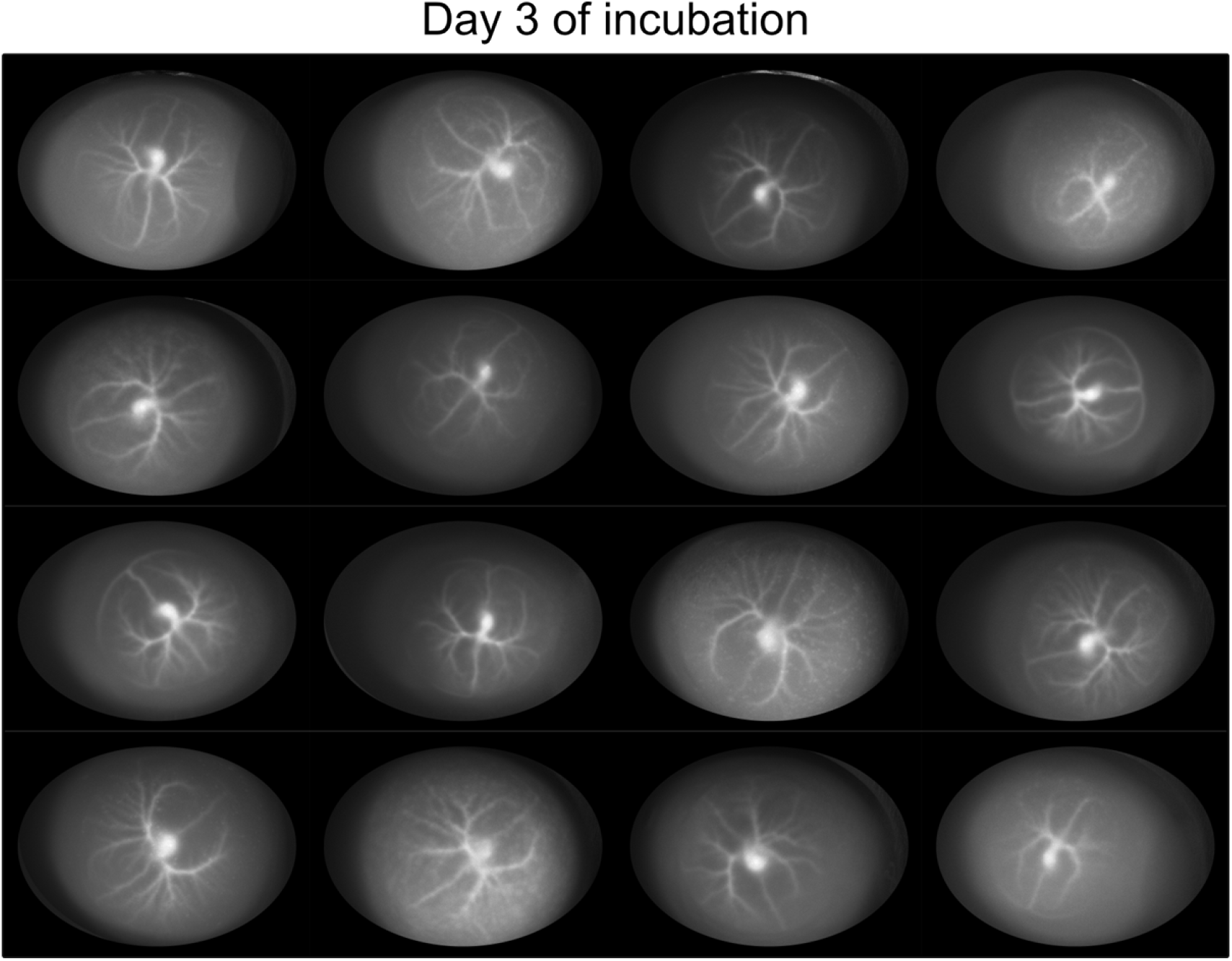
Example of 16 LSCI images before pre-processing. The images were randomly selected from day 3 dataset.

**Figure B2:**
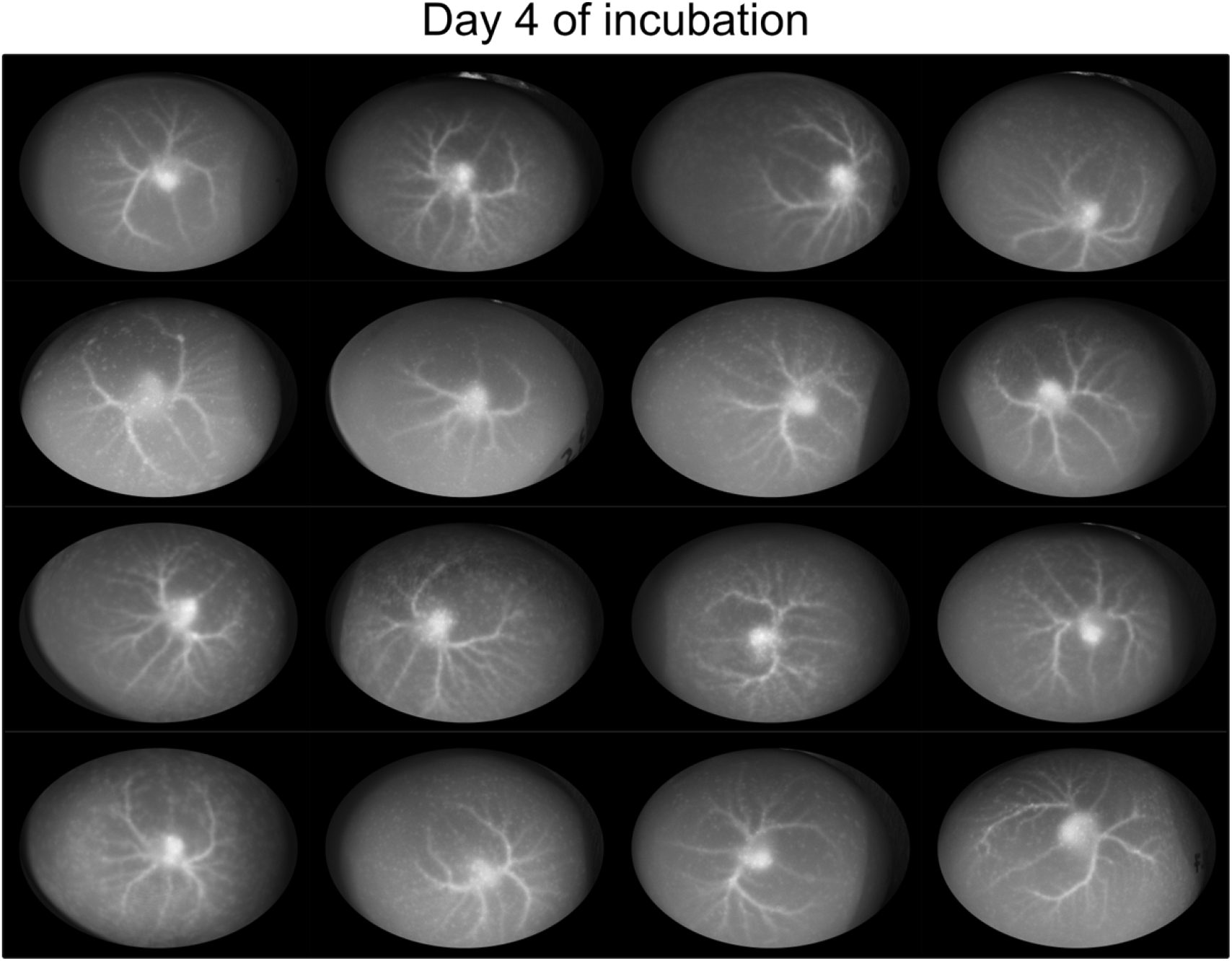
Example of 16 LSCI images before pre-processing. The images were randomly selected from day 4 dataset.

## Data availability

The data that support the findings of this study are available from the corresponding author upon reasonable request.

## Acknowledgements

The authors thanks Caltech students Yusuf Kavranoglu and Weihong (William) Cen for their help and assistance during the deep neural network part of this work. This work was supported by the IST Carver Mead Endowment (25550038).

